# scMagnifier: resolving fine-grained cell subtypes via GRN-informed perturbations and consensus clustering

**DOI:** 10.64898/2026.03.26.714385

**Authors:** Zhenhui He, Kangning Dong

## Abstract

Resolving fine-grained cell subtypes in single-cell RNA sequencing (scRNA-seq) data remains challenging, as their subtle transcriptional differences are often obscured by technical noise and data sparsity. Here, we present scMagnifier, a consensus clustering framework that leverages gene regulatory network (GRN)-informed in silico perturbations to amplify subtle transcriptional differences and uncover latent cell subpopulations. scMagnifier perturbs candidate transcription factors (TFs), propagates perturbation effects through cluster-specific GRNs to simulate post-perturbation expression profiles, and integrates clustering results across multiple perturbations into stable subtype assignments. Additionally, scMagnifier introduces regulatory perturbation consensus UMAP (rpcUMAP), a perturbation-aware visualization that provides clearer separation between cell subtypes and guides the selection of the optimal number of clusters. In both single-batch and multi-batch benchmarks, scMagnifier consistently improves the resolution and accuracy of fine-grained cell type identification. Notably, when integrated with spatial clustering methods such as STAGATE, scMagnifier is compatible with spatial transcriptomics workflows and effectively reveals tumor cell subtypes and their spatial organization in ovarian cancer.

## Introduction

Single-cell RNA sequencing (scRNA-seq) has revolutionized our understanding of cellular heterogeneity by enabling transcriptome-wide profiling at single-cell resolution^1^. Unsupervised clustering is a standard and essential step in the scRNA-seq analysis workflow, routinely used to identify discrete cell types and continuous cell states^2,3^. With the rapid development of clustering algorithms, current tools can consistently and accurately resolve major cell types—such as T and B lymphocytes in immune tissues, neurons and glia in the brain, or epithelial and stromal cells in solid tumors—across a wide range of biological settings. However, resolving fine-grained cell subtypes and accurately delineating their boundaries remains challenging—particularly for transcriptionally similar cell states such as activated versus resting immune cells, malignant epithelial subclones within tumors, or rare cell populations^2,3^. The high dimensionality, sparsity, and noise inherent in scRNA-seq data often obscure subtle but biologically meaningful transcriptional differences, limiting the resolution of clustering methods for fine-grained heterogeneity^4^.

Notably, transcriptionally similar cell populations may respond differently to regulatory perturbations due to variations in their underlying gene regulatory networks (GRNs), offering a powerful means to uncover hidden heterogeneity^5,6^. Several studies have employed in silico gene perturbations to simulate key biological processes, such as cell fate transitions or drug-induced state changes. For example, CellOracle simulates the effects of transcription factor (TF) perturbations on cell identity dynamics by modeling GRN influences, providing interpretable and quantitative insights into developmental trajectories^7^. Similarly, scRank perturbs inferred GRNs to simulate drug-induced regulatory changes, enabling the identification of drug-responsive cell types^8^. In addition, scTenifoldKnk applies GRN perturbation to predict the functional consequences of gene knockouts^9^. Collectively, these studies demonstrate that GRN-informed perturbations provide a powerful approach for revealing functionally relevant cellular heterogeneity. Motivated by these studies, we reasoned that perturbing candidate TFs or genes within GRNs could provide a way to amplify subtle transcriptional differences and thereby facilitate the identification of fine-grained cell subtypes.

Furthermore, different perturbations of regulatory networks can yield distinct patterns of cellular responses, making it necessary to integrate multiple perturbation outcomes to obtain stable clustering results and well-defined subpopulation boundaries. Consensus clustering provides an effective strategy to integrate multiple results and produce stable cluster assignments. Existing consensus clustering approaches typically generate ensemble diversity by repeatedly applying clustering with varying parameters or initializations to the same expression matrix^10-13^, which improves robustness to stochastic variability but fails to enhance the underlying biological signal used to distinguish cell subtypes or states. In contrast, integrating GRN-informed perturbation-derived clustering results into a consensus clustering framework captures the differential responses of subpopulations to distinct regulatory perturbations. This perturbation-aware consensus strategy leads to robust and interpretable boundaries between fine-grained cell types.

Building on these ideas, we develop a GRN-informed perturbation-driven consensus clustering framework named scMagnifier. By integrating with standard clustering algorithms, batch integration methods, or spatial clustering tools, scMagnifier is readily applicable to diverse scenarios—including single-batch, multi-batch, and spatial transcriptomic datasets. Extensive benchmarking demonstrates that scMagnifier consistently improves the resolution and accuracy of fine-grained cell type identification and enhances the detection of rare cell populations. Additionally, scMagnifier integrates perturbation-induced clustering information into the UMAP algorithm, yielding a perturbation-aware visualization, named regulatory perturbation consensus UMAP (rpcUMAP), that provides clearer separation between cell subtypes and guides the selection of the optimal number of cell types.

## Results

### Overview of scMagnifier

The inputs to scMagnifier are the raw gene expression matrix (GEM) and a basic GRN, defined as a set of regulatory interactions from TFs to their target genes (**Fig. 1a**). An initial clustering is obtained using a standard scRNA-seq preprocessing and clustering pipeline implemented in Scanpy^14^. Cluster-specific GRNs are then constructed by pruning the basic GRN based on the expression levels of TFs and their target genes within each cluster (**Fig. 1d and see “Cluster-specific GRN construction” section of the Methods**).

**Fig. 1.**
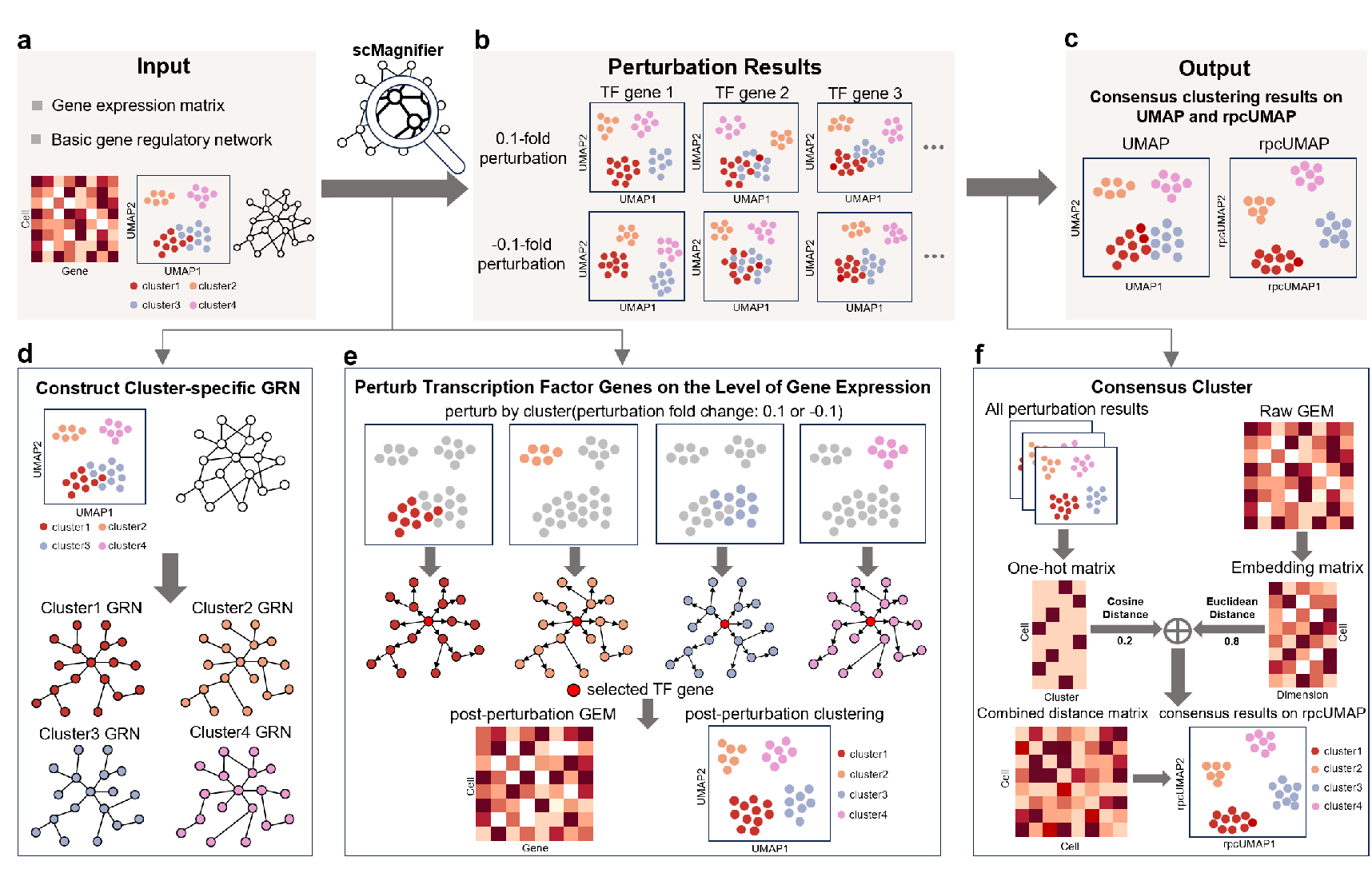
Overview of scMagnifier. **a**. scMagnifier takes the gene expression matrix and a basic gene regulatory network as inputs, and derives the initial clustering results using a standard clustering pipeline. **b**. scMagnifier obtains distinct clustering results by performing TF perturbations on cluster-specific gene regulatory networks. **c**. The output of scMagnifier is a consensus clustering result visualized on UMAP and rpcUMAP embeddings. **d**. scMagnifier first constructs cluster-specific GRN based on the basic GRN and initial clustering results. **e**. scMagnifier systematically perturbs candidate TF genes and propagates their effects through cluster-specific GRN to generate an ensemble of post-perturbation clustering results. **f**. scMagnifier integrates perturbation-driven clustering results with expression-based similarities to construct a combined cell–cell distance matrix, enabling consensus clustering and the generation of a rpcUMAP.

The core of scMagnifier lies in performing GRN-informed in silico perturbations and yielding an ensemble of clustering results across distinct perturbation conditions (**Fig. 1b**). For each candidate TF, a cell-specific perturbation term is first defined relative to its original expression level. This perturbation is then propagated through the corresponding cluster-specific GRN to model downstream regulatory effects (**Fig. 1e and see “TF genes perturbation on the level of gene expression” section of the Methods**). The resulting propagated expression changes are integrated with the original GEM to generate a post-perturbation GEM, from which a perturbation-driven clustering result is obtained.

To derive stable cell subtype assignments, scMagnifier performs consensus clustering on the perturbation-driven ensemble of cluster results (**Fig. 1c, f**). To integrate these clustering results, each clustering outcome is converted into a one-hot matrix, enabling the computation of perturbation-informed cell-cell distances across the ensemble. These perturbation-informed cell-cell distances are further combined with expression-derived distances computed from the embedding matrix of the GEM, resulting in a combined distance matrix that captures both transcriptional similarity and regulatory perturbation-driven differences. Based on this combined distance matrix, a k-nearest neighbor (KNN) graph is constructed for consensus clustering and for generating a regulatory perturbation consensus UMAP (rpcUMAP) (**See “Consensus clustering” section of the Methods**).

To avoid missing subtle but biologically meaningful differences, consensus clustering is initially performed at high clustering resolution. These preliminary clusters are then merged based on inter-cluster centroid distances and a minimum cluster-size threshold to merge closely related or very small clusters into their nearest neighbor clusters and produce the final stable consensus clusters (**Supplementary Fig. S1 and see “Cluster merging” section of the Methods**).

More generally, scMagnifier is extensible and can be applied to multi-batch single-cell datasets. Because both the clustering and the computation of expression-derived distances are performed on low-dimensional embeddings, scMagnifier can directly make use of batch-corrected embeddings (e.g., from Harmony^15^, Scanorama^16^ or scVI^17^), enabling straightforward application to multi-batch datasets (**See “Extension of scMagnifier for multi-batch datasets” section of the Methods**).

### Benchmarking scMagnifier in real datasets

To evaluate scMagnifier’s ability to resolve fine-grained cell subtypes, we benchmarked its performance on multiple publicly available single-cell datasets. For single-batch evaluation, we selected four lung adenocarcinoma datasets that contain rich cellular substructure and published annotations^18^. In each dataset, we fixed the number of output clusters to match the annotation and compared seven methods (Leiden^19^, Louvain^20^, scVI(Leiden)^17^, scVI(Louvain)^17^, SC3s^21^, scMagnifier(Leiden) and scMagnifier(Louvain)) using adjusted Rand index (ARI) and normalized mutual information (NMI) to assess agreement between clustering results and cell type annotations (**See “Benchmarking setup and parameters” section of the Methods**). Across these single-batch benchmarks, scMagnifier consistently achieved the highest ARI and NMI, irrespective of whether Leiden or Louvain was used for clustering, and outperformed a consensus clustering method (SC3s) in our comparisons (**Fig. 2a and Supplementary Table S3, S4**).

**Fig. 2.**
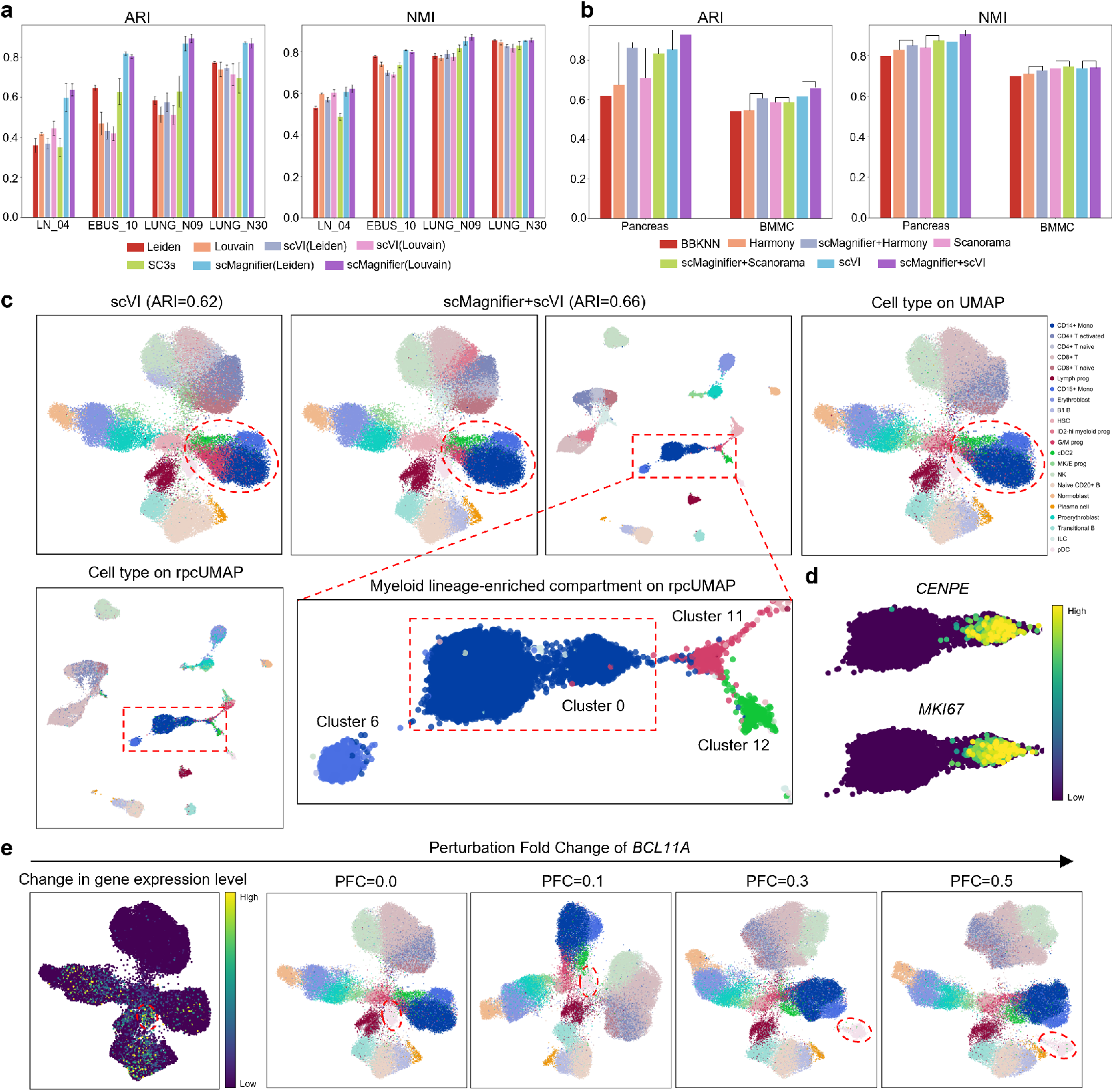
Benchmarking of scMagnifier. **a**. Bar plots illustrating the performance of seven algorithms on four single-batch datasets, evaluated using the Adjusted Rand Index (ARI) and Normalized Mutual Information (NMI). **b**. Bar plots of clustering performance on two multi-batch datasets for seven batch-correction methods, alone or combined with scMagnifier. **c**. UMAP and rpcUMAP visualization of BMMC dataset clustering via scVI (ARI=0.62) and scMagnifier+scVI (ARI=0.66). The red circle highlights a myeloid lineage-enriched compartment containing CD14+ Mono, CD16+ Mono, cDC2 and G/M prog cells. A zoomed view contrasts this compartment on standard UMAP versus rpcUMAP. **d**. rpcUMAP visualization of *CENPE* and *MKI67* gene expression in Cluster 0 of the BMMC dataset, obtained using the scVI+scMagnifier workflow. **e**. UMAP visualization of BCL11A perturbation in the BMMC dataset using the scMagnifier+scVI workflow. The leftmost panel shows the sum of absolute expression changes for *BCL11A* and its target genes across cells. The four panels on the right display UMAP visualizations of post-perturbation clustering results as the perturbation fold change increases.

We next tested scMagnifier’s applicability in multi-batch samples. We benchmarked scMagnifier on two public multi-batch datasets (an integrated pancreas dataset^22^ and the BMMC dataset^23^). For each dataset, we compared several commonly used batch-correction methods alone and in combination with scMagnifier (BBKNN^24^, Harmony^15^, scMagnifier+Harmony, Scanorama^16^, scMagnifier+Scanorama, scVI^17^, scMagnifier+scVI). ARI and NMI were used to quantify agreement with published cell-type annotations. Across both datasets, combining scMagnifier with batch-correction methods resulted in higher ARI and NMI than using batch-correction methods alone (**Fig. 2b and Supplementary Table S5, S6**).

Focusing on the myeloid lineage-enriched compartment in the BMMC dataset, we found that scMagnifier+scVI precisely delineated subcluster boundaries, as validated against reference cell type annotations (**Fig. 2c**). In contrast, scVI drew an inaccurate boundary between granulocyte-monocyte progenitors (G/M prog) and CD14+ monocytes, leading to significant cross-contamination of their identities. Notably, the rpcUMAP visualization generated by scMagnifier+scVI outperformed conventional UMAP by achieving clearer separation of distinct cellular clusters. We further found that rpcUMAP identified two morphologically distinct subpopulations within Cluster 0, which were merged into a single cluster in our benchmarking analyses to align with the predefined cell-type number. Differential gene expression analysis confirmed that these subpopulations exhibited divergent expression of proliferation-associated genes (e.g., *CENPE* and *MKI67*), supporting their classification as distinct subclusters (**Fig. 2d**). Thus, rpcUMAP-based evidence supports revising the optimal number of cell types in this myeloid lineage-enriched compartment from four to five. Collectively, these findings demonstrate that rpcUMAP not only enhances the separation of cell subtypes but also guides the selection of the optimal number of cell types.

Finally, we explored whether scMagnifier amplifies biologically meaningful differences between cells. In the BMMC dataset, using the scMagnifier+scVI workflow, we perturbed the TF gene *BCL11A* and propagated its regulatory effects through cluster-specific GRNs. We observed that plasmacytoid dendritic cell (pDC) clusters exhibited progressively stronger transcriptional changes compared to neighboring cell populations and, as the perturbation fold change increased, gradually dissociated from surrounding clusters (**Fig. 2e**). Importantly, *BCL11A* has been reported as an essential regulator of pDC development and lineage specification^25^. Its knockout leads to impaired pDC development, aberrant lineage specification and a marked decrease in the number and functional maturation of pDCs^25^. These observations demonstrate that gene perturbations can amplify biologically meaningful transcriptional differences, providing a mechanistic explanation for why scMagnifier can resolve fine-grained cell subtypes.

### scMagnifier reveals hidden heterogeneity within MAIT/Th1-Th17 populations

We further evaluate whether scMagnifier can reveal fine-grained cellular heterogeneity that is obscured by conventional clustering. In UPN19_pre dataset^26^, using standard Leiden clustering on the original embedding, Mucosal-associated invariant T (MAIT) cells and a T helper 1/T helper 17 (Th1/Th17)-MAIT mixed population were grouped into a single cluster (Cluster 3) and appeared continuously connected on the original UMAP, indicating limited separability in the original embedding space (**Fig. 3a, b**). Even when the initial resolution was increased, the two cell populations were still recognized as a single cluster (**Supplementary Fig. S3a**). This observation is expected, as MAIT cells exhibit transcriptional programs associated with both Th1 and Th17 effector functions^27^. In contrast, applying scMagnifier to the same cells clearly separated this previously merged population into two distinct clusters (Cluster 2 and Cluster 16) in the rpcUMAP space (**Fig. 3a, b**).

**Fig. 3.**
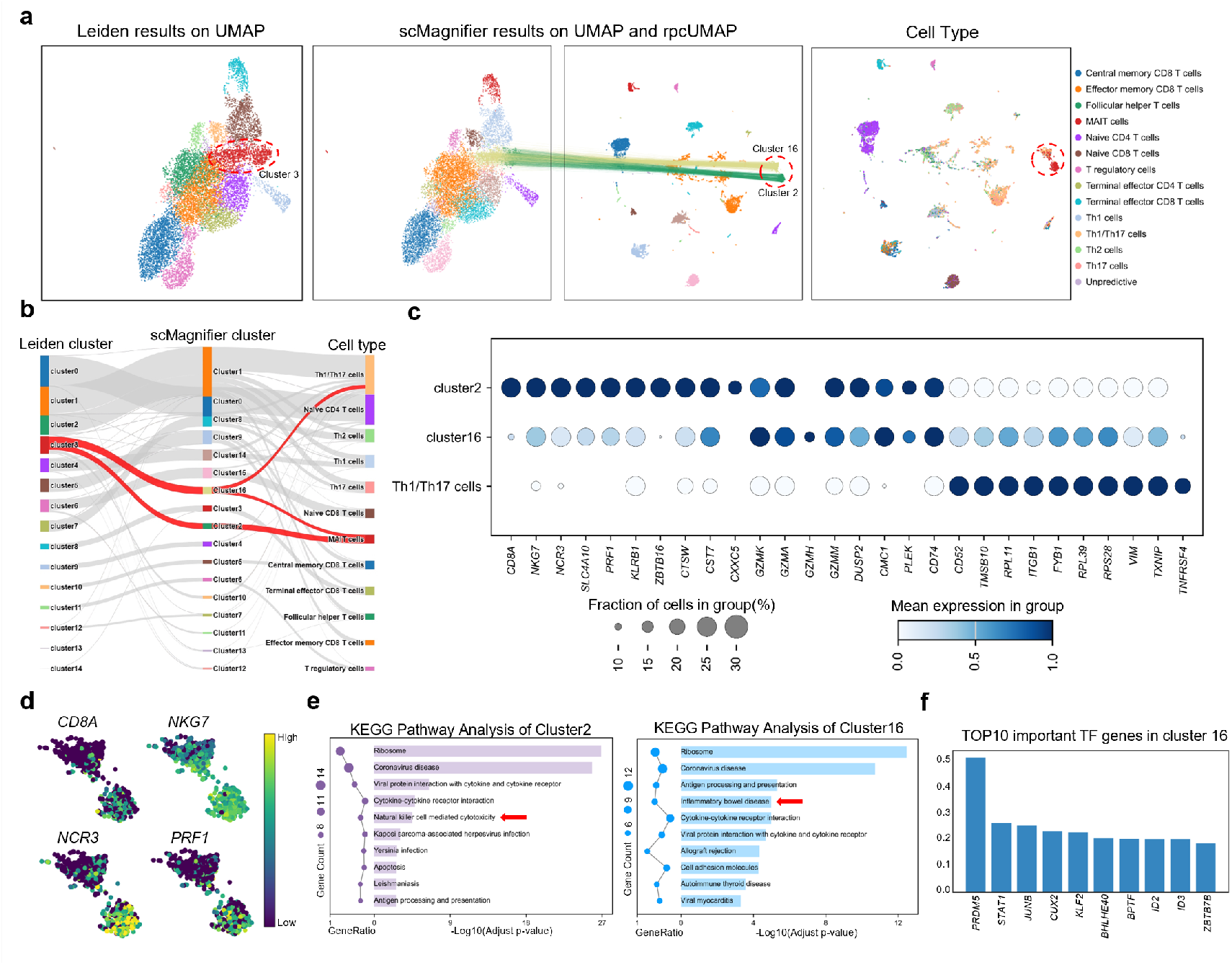
scMagnifier uncovers hidden heterogeneity among MAIT/Th1-Th17 cell populations. **a**. Comparison of original Leiden clustering and scMagnifier results. Left: UMAP visualization of standard Leiden clustering (resolution = 0.75). Middle: UMAP and rpcUMAP visualization of scMagnifier clustering. The connecting lines denote the positional correspondence of clusters identified by scMagnifier between the UMAP and rpcUMAP embedding spaces. The red circle highlights the focal clusters (Cluster 2 and Cluster 16). Right: rpcUMAP visualization of cell type distributions in the dataset. **b**. Sankey diagram showing the correspondence among original Leiden clusters, scMagnifier clusters and cell-type annotations. **c**. Bubble plot of the top 10 differentially expressed genes (DEGs) for each cluster, among cluster 2, cluster 16 and the remaining Th1/Th17 cells. **d**. UMAP heatmaps showing the expression of *CD8A, NKG7, NCR3*, and *PRF1* in cluster 2 and cluster 16. **e**. KEGG pathway enrichment analysis of cluster 2 and cluster 16, showing the top 10 most significantly enriched pathways. **f**. Bar plot of the top 10 TF genes ranked by importance scores in cluster 16.

Differential expression analysis reveals that Cluster 2 is characterized by high expression of cytotoxic-associated genes including *CD8A, NKG7, NCR3* and *PRF1*, while Cluster 16 shows relatively higher expression of several genes associated with Th1/Th17 programs (**Fig. 3c**). Visualization of gene expression on the rpcUMAP provides a more intuitive view of these differences, showing that cytotoxic-associated genes are strongly enriched in Cluster 2 (**Fig. 3d**).

We next performed KEGG pathway enrichment analysis to further validate the functional heterogeneity of these two scMagnifier-resolved clusters (**Fig. 3e**). For Cluster 2, it was significantly enriched in natural killer cell mediated cytotoxicity pathway (Adjusted p-value=3.2×10^-5^), which aligns perfectly with its high expression of cytotoxic genes. Conversely, Cluster 16 was prominently enriched in inflammatory bowel disease pathway (Adjusted p-value=1.03 × 10^-5^), which has been well-documented to be functionally consistent with the molecular features of Th1/Th17 cells^28^.

Meanwhile, we derived the TF importance scores based on the changes induced by the perturbed genes (**See “TF gene importance score calculation” section of the Methods**). We observed *STAT1* ranked as the second most important TF genes in Cluster 16 and it did not appear among the top-ranked TF genes in Cluster 2 (**Fig. 3f and Supplementary Fig. S3b**). *STAT1* has been reported as a key regulator involved in interferon signaling and Th1/Th17-related immune responses^29^.

Collectively, analyses at the gene expression, functional pathway and regulatory levels consistently demonstrate that the two scMagnifier-resolved clusters exhibit biological heterogeneity, which is obscured under conventional clustering but effectively revealed by scMagnifier. Notably, the biological differences observed between the two clusters are consistent with the known plasticity of MAIT cells toward cytotoxic or Th1/Th17-like programs under different contexts^30^. Together, these results highlight scMagnifier’s ability to amplify subtle yet meaningful cellular differences and to facilitate the discovery of fine-grained heterogeneity.

### scMagnifier enables the identification of rare immune cell types with distinct regulatory programs

Rare cells often represent a very small fraction of the total population and are therefore susceptible to technical noise, dropout events and batch effects, while conventional clustering workflows tend to favor dominant transcriptional programs and can merge low-abundance subpopulations into larger clusters^31,32^. To test whether scMagnifier can effectively identify rare cell populations, we applied it to two independent scRNA-seq datasets, EBUS_10 and LUNG_N30^18^. When exploring rare cell states, the minimum cluster-size threshold used during the cluster-merge step can be appropriately reduced based on the size of the dataset and the underlying biological context, so as to avoid merging small but potentially meaningful clusters and to retain candidate rare cell populations for downstream evaluation (**See “ Consensus clustering” section of the Methods**).

In the EBUS_10 dataset, scMagnifier identified two small cell clusters, R1 (18 cells, 0.40%) and R2 (16 cells, 0.36%), that could not be resolved by conventional clustering methods even at high resolution (**Supplementary Fig. S4a**). While these cells were closely positioned within larger populations in the standard UMAP embedding, they became clearly separated from surrounding clusters in the rpcUMAP space (**Fig. 4a**), suggesting the potential presence of subtle but consistent transcriptional differences that are revealed by scMagnifier. To characterize these clusters, we first grouped the remaining cells according to their original cell-type annotations. We then performed differential expression analysis for R1, R2 and each annotation-defined cluster, obtaining cluster-specific differentially expressed gene (DEG) sets for all groups. These cluster-specific DEG sets were subsequently used to quantify transcriptional similarity between R1, R2 and annotated cell populations using the Jaccard coefficient (**See “Jaccard coefficient calculation” section of the Methods**). R1 showed similarity with germinal centers (GC) B cells in the dark zone (DZ) (Jaccard = 0.33) and lower similarity with mucosa-associated lymphoid tissue (MALT) B cells (Jaccard = 0.12), while R2 exhibited a similarity with GC B cells in the DZ (Jaccard = 0.41) (**Supplementary Fig. S4b and Supplementary Table S7**). Consistent with these results, violin plots of DEG expression (**Fig. 4b**) revealed that R1 shares partial expression patterns with both MALT B cells and GC B cells in the DZ, while also displaying distinct expression distributions. In contrast, R2 more closely resembled GC B cells in the DZ but still exhibited differences. Notably, cells in R1 and R2 were originally annotated as MALT B cells and GC B cells in the DZ (**Fig. 4a**). Together, these results suggest that R1 likely represents a finer-grained subpopulation within the MALT B cell compartment, and R2 likely represents a finer-grained subpopulation within GC B cells in the DZ.

**Fig. 4.**
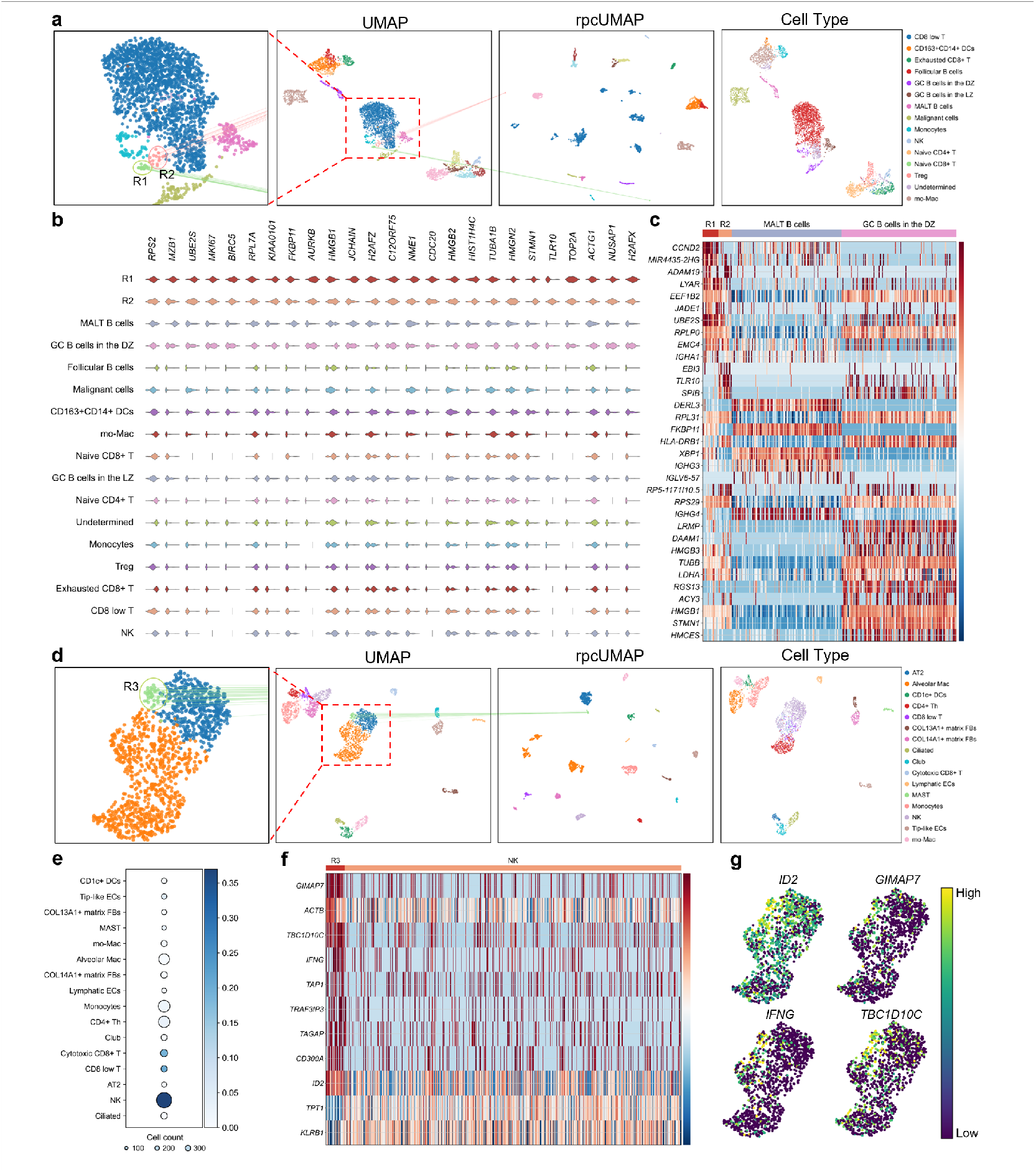
scMagnifier detects rare immune cell populations in lung adenocarcinoma datasets. **a**. UMAP and rpcUMAP visualizations of scMagnifier results on the EBUS_10 dataset, highlighting the R1 and R2 clusters. The connecting lines denote the positional correspondence of R1 and R2 between the UMAP and rpcUMAP embedding spaces. **b**. Violin plots showing the expression profiles of the top 15 DEGs individually selected from R1 and R2, across R1, R2 and the remaining cell-type-annotated cells. **c**. Heatmap of gene expression for the top 10 upregulated genes selected from each of the four clusters (R1, R2, MALT B cells and GC B cells in the DZ). **d**. UMAP and rpcUMAP visualizations of scMagnifier results on the LUNG_N30 dataset, highlighting the R3 clusters. **e**. Bubble plot showing Jaccard coefficient values between R1 and clusters of the remaining cells grouped by cell-type annotations. **f**. Heatmap of gene expression for the top 10 upregulated genes selected from each of the two clusters (R3 and NK). **g**. UMAP heatmaps showing the expression of *ID2, IFNG, GIMAP7* and *TBC1D10C* in R3 and NK cells.

To further examine the transcriptional differences among these populations, we next visualized the significantly upregulated genes for R1, R2, MALT B cells and GC B cells in the DZ using a heatmap (**Fig. 4c**). R1 exhibits significantly higher expression of *CCND2* and *UBE2S* relative to the other cell clusters. It has been reported that *CCND2* is implicated in promoting B cell cell-cycle entry and has been linked to proliferative B cell states^33^, while *UBE2S* contributes to mitotic progression and supports proliferation in multiple cellular systems^34^. Together, these findings suggest that R1 represent a rare subpopulation of MALT B cells associated with proliferation and activation. By contrast, R2 shows higher expression of *EBI3* and *TLR10* relative to the other cell clusters. *EBI3* has been reported in activated B cells and can modulate B cell differentiation and immune responses^35^, and *TLR10* has been associated with activation and mucosal immune regulation^36^. These observations suggest that R2 represents a rare subpopulation of B cells associated with activation and immune regulation, likely resembling a MALT-like subcluster rather than a simple proliferative GC-DZ program.

Similarly, we next applied scMagnifier to the LUNG_N30 dataset and identified a small cluster, R3 (36 cells, 1.25%), that was not detected by conventional high-resolution clustering (**Supplementary Fig. S4a**). In the standard UMAP embedding, R3 appeared connected to neighboring cells, whereas in rpcUMAP it formed a clearly separated cluster (**Fig. 4d**). Jaccard similarity between R3 DEGs and annotated clusters reveals highest overlap with natural killer (NK) cells (Jaccard = 0.37) (**Fig. 4e and Supplementary Table S8**), and R3 cells were originally annotated as NK (**Fig. 4d**), supporting the hypothesis that R3 is a rare NK subpopulation. To investigate this further, we compared significantly upregulated genes for R3 and NK cells (**Fig. 4f**). R3 exhibits significantly higher expression of *ID2, IFNG, GIMAP7* and *TBC1D10C* relative to NK cell clusters, which is also clearly reflected in the UMAP visualization (**Fig. 4g**). It has been reported that *ID2* is a key transcription factor regulating NK cell development and maturation^37^, while *IFNG* is a hallmark effector cytokine of NK cells^38^. *GIMAP7* and *TBC1D10C* are involved in lymphocyte activation and regulation^39,40^. Together, these findings suggest that R3 represents a rare subpopulation of NK cells associated with a distinct activation or maturation state, potentially corresponding to the CD56^bright^ NK cells, which are known to express higher levels of *IFNG*, and are often linked to early stages of activation and immune responses^41^.

In summary, scMagnifier successfully identified rare subpopulations in the EBUS_10 and LUNG_N30 datasets, highlighting its potential to uncover biologically meaningful yet low-abundance cell populations that are often overlooked by conventional clustering methods.

### Integration of scMagnifier with STAGATE reveals tumor cell subtypes and spatial organization in ovarian cancer

The identification of tumor cell subtypes and their spatial organization plays a critical role in understanding cancer progression and therapeutic response^42^. STAGATE enables the accurate identification of spatial domains by learning low-dimensional latent embeddings^43^, while scMagnifier provides fine-grained types of cells. Hence, we integrated scMagnifier with STAGATE (**See “Integration of scMagnifier with STAGATE for Spatial Transcriptomics Analysis” section of the Methods**), aiming to identify biologically meaningful tumor subtypes and illustrate their spatial domains. We used the spatial transcriptomics dataset of epithelial ovarian cancer from the SPATCH website^44^, which incorporates both detailed cell annotations and spatial layer annotations. For our analysis, we selected a subset of cells located at the center of spatial coordinates comprising 48,793 cells (**Fig. 5b**).

**Fig. 5.**
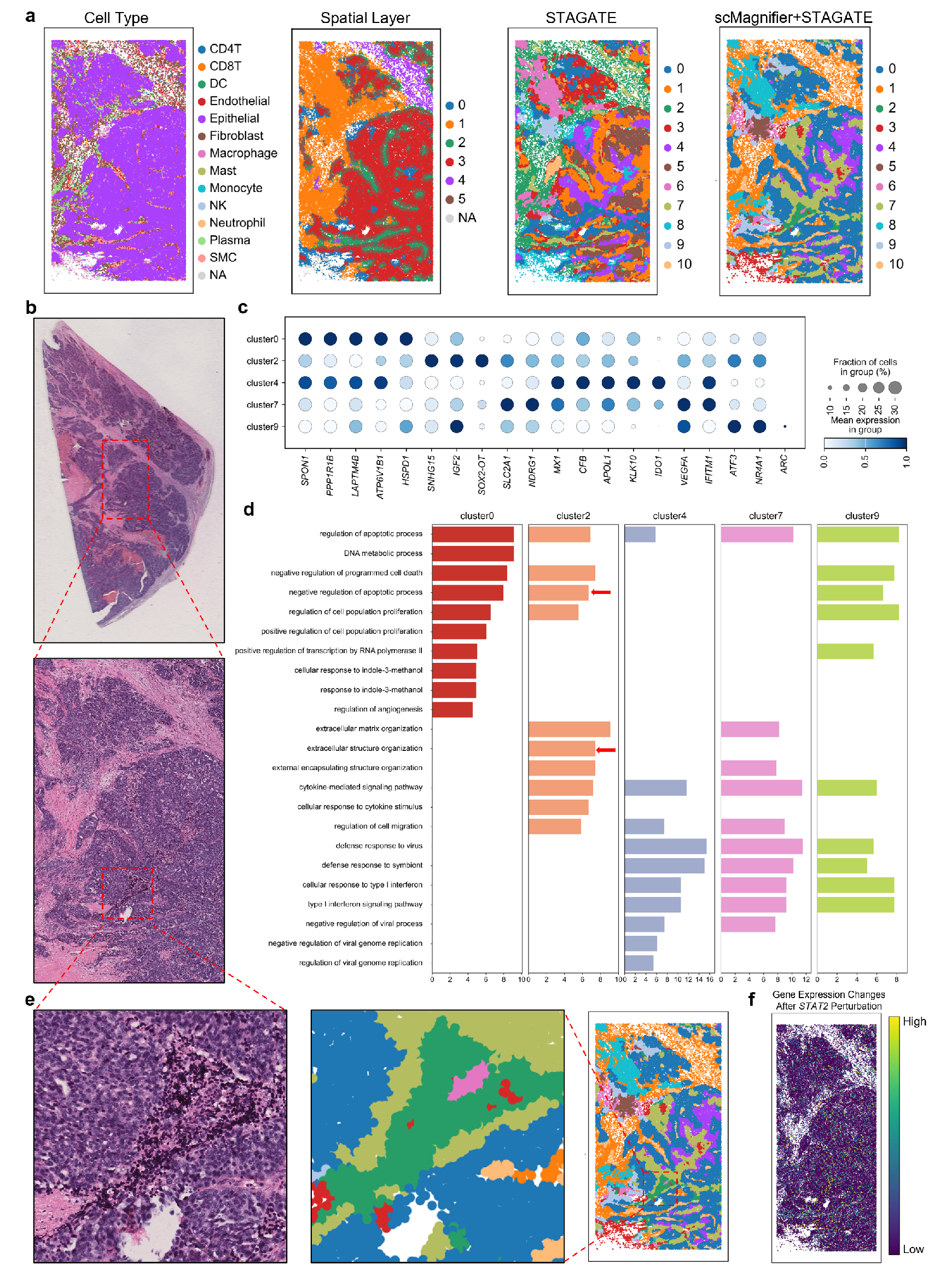
The integration of scMagnifier and STAGATE identifies tumor cell subtypes and their spatial organization in ovarian cancer. **a**. Spatial visualization of the ovarian cancer dataset. Left two panels: cell-type annotations and spatial layer annotations provided with the dataset. Right two panels: clustering results from STAGATE alone (Leiden, resolution=0.3) and from scMagnifier combined with STAGATE. **b**. H&E-stained image highlighting the specific regions analyzed in the dataset used for our experiments. **c**. Bubble plot of the top 5 differentially expressed genes for each of the five tumor subclusters identified by scMagnifier+STAGATE. **d**. GO enrichment analysis of the five tumor subclusters, showing the top 10 significantly enriched GO terms for each subcluster. **e**. Spatial correspondence between histology and scMagnifier+STAGATE clustering. Left panel: dark regions highlighted in the H&E-stained image. Right panel: spatial map showing the location of cluster 2 identified by scMagnifier+STAGATE. **f**. Spatial heatmap showing the sum of absolute expression changes for *STAT2* and its target genes across cells after *STAT2* perturbation.

Epithelial ovarian cancer, a malignancy originating from epithelial cells, is characterized by significant cellular heterogeneity and complex tumor microenvironment^45^. Based on cell annotations and spatial layer annotations provided by the dataset (**Fig. 5a**), we determined the approximate location of the tumor’s core regions (The overlapping region of Epithelial cell clusters in the cell type annotation map and Layer 3 regions in the spatial layer annotation map). Subsequently, using the final clustering results from scMagnifier combined with STAGATE, we identified five distinct subclusters of tumor cells (Cluster 0, Cluster 2, Cluster 4, Cluster 7 and Cluster 9), which correspond to potential tumor core regions (**Fig. 5a, b**). Using differential gene expression analysis and functional enrichment analysis, we confirmed that the five tumor subclusters exhibit distinct molecular and functional differences (**Fig. 5c, d**).

Notably, we observed that the spatial localization of Cluster 2 closely overlapped with the deeply stained regions in H&E histology (**Fig. 5e**). Differential gene expression analysis revealed that Cluster 2 displayed high expression of *IGF2* (**Fig. 5c**). It has been reported *IGF2* is involved in tumor growth and invasion mechanisms^46^. Consistent with this, functional enrichment analysis identified pathways related to negative regulation of apoptosis (Adjusted p-value=2.0×10^-7^) and extracellular structure organization (Adjusted p-value=3.9×10^-8^) in Cluster 2 (**Fig. 5d**), indicating that these cells may actively evade cell death and promote metastatic potential. This functional profile aligns well with *IGF2*-mediated proliferative and anti-apoptotic signaling, a characteristic feature of aggressive epithelial tumor subpopulations^46^. Importantly, deep staining in H&E images generally indicates high cellular density and malignancy, which are hallmarks of invasive tumor subpopulations^47^, and further supports that Cluster 2 corresponds to a highly aggressive tumor subpopulation. Collectively, this spatial correlation suggests that, by integrating scMagnifier with STAGATE, we were able to identify high-invasion tumor regions without relying on traditional histological images, a capability that was not achievable with STAGATE alone.

To further investigate why this specific region was detected, we analyzed the impact of TF genes perturbation within the scMagnifier framework. After perturbing *STAT2*, we calculated the changes in gene expression levels of *STAT2* and its target genes, and then visualized the resulting changes on the spatial map (**Fig. 5f**). Notably, the region corresponding to cluster 2 became significantly more prominent compared to its surrounding areas, suggesting that this cluster was more strongly affected by *STAT2* perturbation. This increased prominence allowed for the successful identification of this region after *STAT2* perturbation (**Supplementary Fig. S5**), indicating that perturbation amplified the underlying biological differences within this cluster. In summary, the integration of scMagnifier with STAGATE enables the revelation of tumor cell subtypes and spatial organization.

## Discussion

Accurately delineating cell subpopulations and their boundaries remains challenging, particularly when transcriptional differences are subtle or obscured by noise, batch effects. In this study, we presented scMagnifier, a regulatory perturbation-driven consensus clustering framework designed to resolve fine-grained cell subtypes. scMagnifier first amplifies subtle regulatory differences through systematic perturbation of TF genes to reveal latent subclusters, and then integrates clustering outcomes across multiple perturbations via consensus clustering to obtain stable subtype assignments and well-defined boundaries. We demonstrated practical utility of scMagnifier in several applications. We found scMagnifier clearly reveals hidden heterogeneity within MAIT/Th1-Th17 populations. We additionally substantiated the ability of scMagnifier to detect rare cell populations. Finally, by integrating scMagnifier and STAGATE, we identified five subtypes of ovarian cancer and detected invasive regions without incorporating pathological images.

Recent studies have demonstrated that GRN perturbation models are capable of uncovering latent biological signals^7-9^. Building on this framework, we showed that perturbing TF genes at the gene expression level amplifies the underlying biological differences, thereby enabling the identification of subclusters (**Fig. 2e and Fig. 5f**). It is precisely this perturbation strategy that supports the key advantage of scMagnifier. However, our perturbations do not explicitly model true post-perturbation cellular states, as they rely on GRN-based signal propagation at the transcriptomic level rather than learning distributional shifts induced by real perturbations. To address this limitation and enhance biological fidelity, future extensions could integrate scMagnifier with optimal transport-based perturbation models such as CellOT^48^, which learn mappings between control and perturbed single-cell distributions, thereby improving the biological realism of perturbation effects while retaining scMagnifier’s ability to amplify regulatory differences for fine-grained subcluster discovery.

scMagnifier effectively identifies heterogeneous cell populations, which form clearly separated clusters from the remaining cells in the rpcUMAP embedding (**Fig 3a and Fig. 4a, d**). This clear separation stems from two synergistic factors: first, the perturbation of TF genes amplifies the underlying regulation-informed transcriptional differences between cell subpopulations; second, unlike standard UMAP, rpcUMAP uniquely incorporates perturbation-derived intercellular distance information. By integrating this perturbation-based metric, rpcUMAP further exaggerates the dissimilarities between distinct clusters, thereby achieving the enhanced separation observed in the embedding. However, many cell states are defined by features beyond transcript abundance (such as protein markers, chromatin accessibility) and therefore may remain ambiguous in transcription-only analyses. Extending scMagnifier to integrate multimodal measurements via modality-aware distance fusion or multimodal GRN inference would increase sensitivity and specificity for cell type detection.

## Methods

### Data preprocessing

All datasets were prepared in the AnnData format and processed using Scanpy^14^. First, lowly expressed genes were filtered out by retaining only genes with at least one count across all cells. Expression values were then normalized per cell to a total UMI count of 10,000. Highly variable genes (HVGs) were identified on the normalized non-log-transformed expression matrix, and the top 2,000 HVGs were retained. After subsetting the dataset to these HVGs, expression values were renormalized per cell to a total UMI count of 10,000 to account for the effects of gene filtering. Importantly, the renormalized non-log-transformed GEM was retained and used for downstream cluster-specific GRN construction and TF gene perturbation analyses. Subsequently, the expression values were log-transformed and scaled to a maximum value of 10. Principal component analysis was performed with 20 components, followed by the construction of a KNN graph (k=10) and UMAP for dimensionality reduction. Finally, initial cell clustering results were obtained using the Leiden^19^ or Louvain^20^ algorithm.

### Cluster-specific GRN construction

Cluster-specific GRNs were constructed following the standard CellOracle framework^7^. As prior regulatory information, we employed the human promoter-based GRN provided by CellOracle as our basic GRN, which defines potential TF gene-target gene relationships. Cluster-specific GRNs were constructed based on the basic GRN and initial clustering results, using the renormalized but non-log-transformed GEM stored during data preprocessing. Firstly, dimensionality reduction was performed using PCA (oracle.perform_PCA() function), followed by KNN-based imputation (oracle.knn_imputation() function) to alleviate data sparsity. Regulatory relationships between TF genes and their putative target genes were modeled using regression-based approaches (oracle.get_links() function), and cluster-specific regulatory links were further identified and filtered based on statistical significance and network scores (links.filter_links() function). Finally, quantitative GRN models for perturbation simulation were fitted (oracle.fit_GRN_for_simulation() function). All parameters were set according to the default or recommended settings in CellOracle.

### TF genes perturbation on the level of gene expression

TF gene perturbation was performed on the renormalized but non-log-transformed GEM retained during data preprocessing (**X** ∈ ***R***^*N×G*^). This design ensured that perturbations were applied in the original linear expression space, thereby being consistent with the linear regulatory assumptions underlying the GRN models, which were inferred from the same renormalized non-log-transformed matrix.

Candidate TF genes for perturbation were selected by intersecting the TF genes from the basic GRN with the top 2,000 HVGs. Perturbations were applied in a cluster-wise manner: let the dataset be partitioned into *C* clusters according to the initial cluster results and let *I*_*c*_ denote the set of cell indices belonging to cluster c. For a selected perturbed TF gene *g*_0_ and perturbation fold change *μ* (*μ* ∈{−0.1, 0.1}), the initial, cluster-specific perturbation matrix **ΔX**^(0,*c*)^ ∈ ***R***^*N*×*G*^was defined by

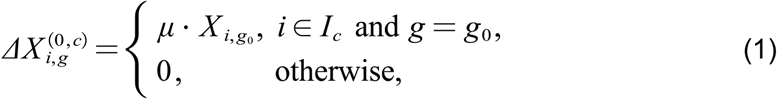

where ***X***_*i,g*_ denotes the original expression of gene *g* in cell *i*.

For each cluster *c*, let **C**^(*c*)^ ∈ ***R***^*G*×*G*^ denote the cluster-specific regulatory coefficient matrix. Propagation of the initial perturbation through the cluster-specific GRN was performed iteratively to capture multi-step regulatory effects. For iteration *k* = 1, …, *n* (we use *n* = 3 iterations), we computed

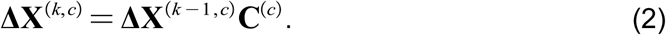

To preserve the intended perturbation on the TF itself across iterations, the perturbed-TF column was restored to its initial values before proceeding to the next iteration.

After *n* iterations, the propagated perturbation for cluster *c* was **ΔX**^(*n,c*)^. The overall propagated perturbation affecting all cells was obtained by summing cluster-wise contributions:

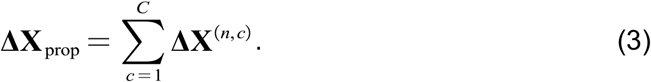

The post-perturbation GEM was then formed by adding the propagated perturbation back to the original expression matrix:

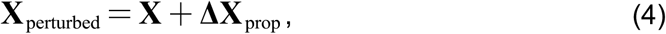

and negative values were clipped to zero to maintain non-negativity. Finally, **X**_perturbed_ was processed through the downstream analytical pipeline consistent with that used in our data preprocessing to produce the post-perturbation clustering.

This procedure was repeated independently for each perturbation candidate TF gene producing a collection of clustering outcomes that together formed the perturbation-driven ensemble.

### Consensus clustering

To integrate perturbation-induced clustering results and generate a consensus, we first transformed the clustering assignments into a one-hot matrix. For each clustering result, we assigned a “1” to the cluster a cell belongs to and “0” to other clusters, producing a sparse matrix for each clustering result. These one-hot matrices were then concatenated to form a high-dimensional matrix where each row represents a cell, and each column corresponds to a particular cluster assignment across all perturbation experiments. Next, we computed the pairwise cosine distance between cells based on this one-hot matrix (**Supplementary Fig. S7**). The cosine distance between two cells *i* and *j*, was calculated as

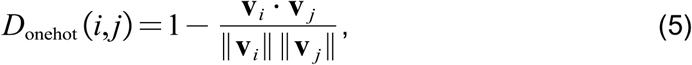

where **v**_*i*_ and **v**_*j*_ are the one-hot vectors for cells *i* and *j*.

In parallel, we calculated the Euclidean distance between cells in the embedding space (PCA-reduced space in single-batch scRNA-seq datasets) derived from the original GEM. The Euclidean distance between cells *i* and *j* in the *k*-dimensional (*k* = 20) embedding space was given by

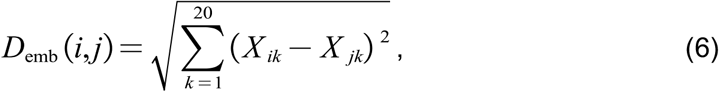

where *X*_*ik*_ and *X*_*ik*_ are the *k*-th embedding values of cells *i* and *j*.

Both the one-hot distance matrix and embedding distance matrix were min-max normalized to ensure their scales were consistent by

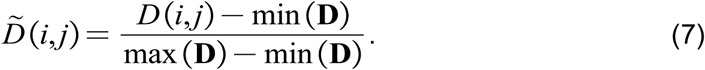

Then we computed a weighted sum of the normalized one-hot and embedding distances by

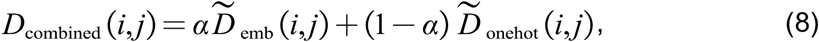

where *α* is set as 0.8 by default (**Supplementary Fig. S6**).

We next computed rpcUMAP with the combined distance matrix as the precomputed distance metric (via umap.umap_.UMAP() function) and set the number of nearest neighbors to 10. We further constructed a KNN graph (k=10) from the combined distance matrix and applied the clustering algorithm (consistent with the clustering approach implemented in our data processing), with the resolution parameter specified as 1.5.

### Cluster merging

We performed cluster merging to produce the final stable consensus clusters. This process included two stages: centroid-based merging and merging of small clusters.

In the first stage, we calculated the centroids of each cluster. The centroid of a cluster was computed from the HVG expression matrix, which corresponded to the final output of our preprocessing pipeline, where gene expression values had already been normalized, log-transformed and scaled. Specifically, the centroid of a cluster *l* was defined as the arithmetic mean of the HVG expression vectors of all cells assigned to that cluster:

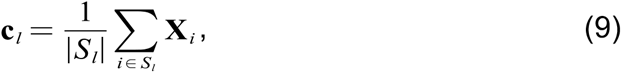

where *S*_*l*_ represents the set of cells assigned to cluster *l*, |*S*_*l*_| is the number of cells in cluster *l* and **X**_*i*_ denotes the HVG expression vector of cell *i*.

The centroids were then used to calculate the pairwise Euclidean distances between them:

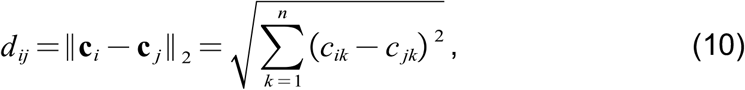

where *c*_*ik*_ and *c*_*jk*_ represent the expression values of gene *k* in the centroids ***c***_*i*_ and ***c***_*j*_ respectively, and *n* is the number of HVGs. Then we computed the median of the minimum nearest-neighbor distances for each cluster, and multiplied this value by a scaling factor (set as 0.75 by default, **Supplementary Fig. S8**) to obtain the final distance threshold for merging similar clusters:

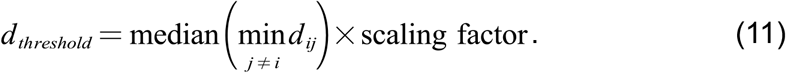

If the centroid distance between two clusters was smaller than this threshold, the clusters were merged.

In the second stage, we merged small clusters. Small clusters were defined as those containing fewer cells than a minimum threshold, set as a fraction of the total number of cells (default: 1% of total cells). For each small cluster, the nearest non-small cluster was identified based on the centroid distance, and the small cluster was merged with this centroid-nearest non-small cluster. This step was originally intended to merge boundary-associated cells misidentified as small clusters. However, if the algorithm is applied to the task of identifying rare cell populations, the threshold can be reduced (e.g., set 0.1% of total cells) to retain low-abundance cell populations. Notably, adjusting this threshold also enables flexible control over the final number of clusters obtained after merging.

### Extension of scMagnifier for multi-batch datasets

scMagnifier can be adapted to handle multi-batch datasets by integrating it with batch effect correction methods that operate in the dimensionality reduction space, such as Harmony^15^, Scanorama^16^ or scVI^17^. Specifically, during the algorithm workflow, we make two key adjustments. First, during data preprocessing and TF genes perturbation, we replace PCA-based dimensionality reduction with a batch effect correction method. Second, in the consensus clustering step, we use the embedding obtained from the batch effect correction method to compute cell distances in the reduced space, instead of using the PCA matrix. All other steps remain unchanged.

### Benchmarking setup and parameters

In our benchmarks, we ensured that the number of output clusters was kept consistent with the annotated cell-type count. Specifically, during the data preprocessing step, we adjusted the clustering resolution to match the number of clusters in the cell annotations. This resolution was maintained throughout the TF genes perturbation process. Finally, during the cluster merging stage, we controlled the final number of clusters by adjusting the minimum threshold to ensure it aligned with the annotated cell-type count. All other parameters across algorithms were kept at their default settings.

### Identifying differentially expressed genes

We used the Wilcoxon test implemented in SCANPY^14^ to identify DEGs. For general analyses, DEGs were selected based on the ranking scores generated by the test combined with an adjusted p-value (Benjamin-Hochberg-corrected FDR) < 0.05. When analyzing whether rare cells constitute distinct subtypes, we selected strictly upregulated differentially expressed genes (DEGs) using dual thresholds: adjusted p-value (Benjamin-Hochberg-corrected FDR) < 0.05 and log fold change (logFC) > 0.25 (**Fig. 4c, f**).

### Pathway enrichment analysis

We first used the Wilcoxon test implemented in SCANPY^14^ to identify DEGs. For each cluster, DEGs were sorted by adjusted p-value (ascending) and logFC (descending), and the top 200 DEGs were selected for enrichment analysis. Subsequently over-representation analysis (ORA) was conducted using the enrichr function in the GSEApy^49^ package for two functional databases: GO Biological Process 2021 and KEGG 2021 (Human), with a significance threshold of adjusted p-value < 0.05.

### TF gene importance score calculation

To quantify how strongly each perturbed TF influences a given consensus cluster, we computed a per-gene, per-cluster importance score. Let *G* be the set of candidate TF genes and let *I* be the set of cells. For a perturbation of gene *g* ∈ *G*, we denoted post-perturbation GEM by **X**^(*g*)^ ∈ *R*^|*I*|×*P*^(*P* is the number of genes) and the original (renormalized but non-log-transformed) GEM by **X**.

We first calculated a binary change indicator to quantify perturbation-induced cluster assignment shifts for each cell. For each perturbation gene *g*, we first established a label mapping between original and perturbed clusters to account for potential relabeling of identical clusters. Let *L*_*i*_ be the original label of cell *i* and 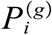 the label of cell *i* after perturbing gene *g*. We defined the overlap matrix **M**^(*g*)^:

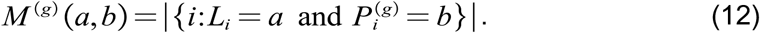

We obtained a mapping ***m***^(*g*)^ from each original cluster *a* to a perturbed cluster by choosing the perturbed label with maximal overlap:

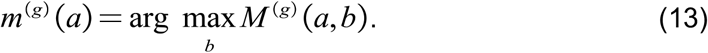

Using this mapping, we defined the per-cell binary change indicator

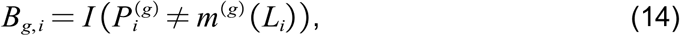

where *I*(·) is the indicator function. The binary matrix **B** ∈{0,1}^|*G*|×|*I*|^ collected *B*_*g,i*_.

We then computed a continuous change metric to quantify the magnitude of global expression profile alterations for each cell under perturbation *g*. Let *μ*_*j*_ and be *σ*_*j*_ the mean and standard deviation (across cells) of log(1 +*X*_:,*j*_) for gene *j*, computed from the original GEM **X**. We defined

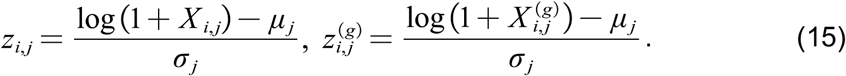

The continuous change for cell *i* under perturbation *g* was

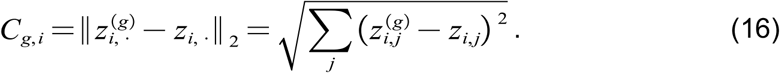

The continuous matrix **C** ∈ *R*^|*G*| ×|*I*|^ collected *C*_*g,i*_.

To combine the binary and continuous signals on a common scale, each row **C**_*g*_ was min-max normalized across cells:

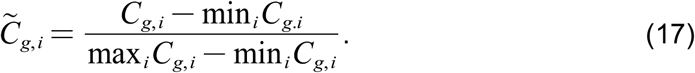

The two signals were then combined to yield a per-gene per-cell combined perturbation score

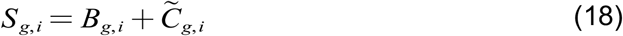

Given consensus cluster *k* with cell set *I*_*k*_, the importance of TF gene *g* for cluster *k* is the mean of perturbation scores across cells in that cluster:

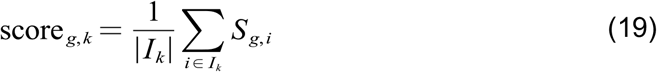

### Jaccard coefficient calculation

To assess the transcriptional similarity between the rare cell clusters and biologically annotated cell populations, we quantified the overlap of their cluster-specific DEG sets using the Jaccard similarity coefficient. For each cluster, cluster-specific DEGs were first identified and the top 50 DEGs were selected to form distinct gene sets for each cluster. The Jaccard coefficient between the gene set of a rare cell cluster *G*_1_ and the gene set of an annotated cell population *G*_2_ was calculated using the following formula:

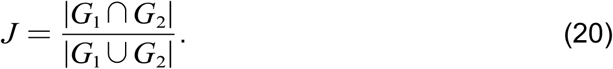

### Integration of scMagnifier with STAGATE for Spatial Transcriptomics Analysis

To integrate scMagnifier with STAGATE^43^ for spatial transcriptomics analysis, we followed the same extension strategy as described for applying scMagnifier to multi-batch datasets. Specifically, STAGATE was used to generate a low-dimensional embedding that captures spatial context, and this embedding replaced the PCA space in the scMagnifier workflow. During data preprocessing and TF gene perturbation, the STAGATE-derived latent embedding was used for downstream analyses instead of PCA. In the consensus clustering step, cell-cell distances in the reduced space were computed based on the STAGATE embedding, while all other steps of the scMagnifier algorithm remained unchanged.

In our dataset, Spatial neighbor graphs in STAGATE were constructed using a KNN-based strategy (*k*=10). STAGATE was trained for 300 epochs to obtain stable spatial embeddings. Downstream clustering was performed using the Leiden algorithm with a resolution of 0.3.

## Data availability

All data analyzed in this paper are available in raw form from their original authors. Specifically, the lung adenocarcinoma datasets are available at NCBI Gene Expression Omnibus (GEO) (https://www.ncbi.nlm.nih.gov/geo/query/acc.cgi?acc=GSE131907). The pancreas dataset is collected from NCBI GEO (https://www.ncbi.nlm.nih.gov/geo/query/acc.cgi?acc=GSE84133). The BMMC datasets is available at NCBI GEO (https://www.ncbi.nlm.nih.gov/geo/query/acc.cgi?acc=GSE194122). The UPN19_pre dataset is accessible on Human Antigen Receptor Database (huARdb) website (https://huarc.net/v2/database/browse/UPN19_pre). The ovarian cancer dataset and H&E images are available at SPAtial Transcriptomics resource for subCellular and High-throughput platforms (SPATCH) website (https://spatch.pku-genomics.org/#/dataset/xenium).

## Code availability

The scMagnifier algorithm is implemented in Python and is available on Github (https://github.com/RucDongLab/scMagnifier).

